# Sex-Specific Plasticity Explains Genetic Variation in Sexual Size Dimorphism in *Drosophila*

**DOI:** 10.1101/2021.06.16.448738

**Authors:** Isabelle M Vea, Austin Wilcox, W. Anthony Frankino, Alexander W Shingleton

## Abstract

The difference in body size between females and males, or *sexual size dimorphism* (SSD), is almost ubiquitous, and yet we have a remarkably poor understanding of the developmental-genetic mechanisms that generate it. Such an understanding is important if we are to distinguish between the many theoretical models of SSD evolution. One such model is the condition dependence hypothesis, which proposes that the body size of the larger sex is also more environmentally sensitive, a phenomenon called *sex-specific plasticity* (SSP). Because SSP generates differences in female and male body size, selection on plasticity may underlie the evolution of sexual size dimorphism. To test this hypothesis, however, we need to know the genetic architecture of both SSD and SSP, which is challenging because both are characteristics of populations not individuals. Here, we overcome this challenge by using isogenic lineages of *Drosophila* to measure both SSD and SSP for a genotype. We demonstrate extensive genetic variation for SSD among genotypes that is tightly correlated with variation in SSP, indicating that the same developmental-genetic mechanisms regulate both phenomena. These data support the condition dependence hypothesis and suggest that the observed SSD is a consequence of selection on the developmental-genetic mechanisms that regulate SSP.

## Introduction

Sexual Size Dimorphism (SSD), the difference in body size between males and females, is perhaps the most familiar and widespread form of sexual dimorphism. SSD is extremely variable among species regardless of their average size, and varies in both direction and magnitude [1,2]. For example, most mammals and birds have male-biased SSD, where males are larger than females [3,4], whereas insects typically exhibit female-biased SSD [5]. SSD is also highly evolutionarily labile, even among closely related species and populations within species [6–9] suggesting that there is considerable intraspecific genetic variation in SSD upon which selection can act. While numerous hypotheses have been proposed regarding the selective pressures that might give rise to SSD [10], the developmental-genetic mechanisms that are the target of these selective agents remain largely unknown. This represents a substantial gap in our understanding of the evolution of SSD, since knowing the targets of selection could provide insight regarding the modes of selection favoring sex-specific changes in body size. Hampering efforts to fill this gap is the challenge of describing genetic variation in SSD within a population, since SSD cannot be measured for an individual.

For SSD to evolve, the strength or direction of selection on male and female body size must necessarily differ. For example, sex-specific selection on body size can produce male-biased SSD when focused on display or fighting ability, whereas female-biased SSD can result from fecundity selection, which also typically correlates with body size. Regardless of the ultimate selective pressures that generate SSD, theory suggests that the larger sex gains greater fitness benefits from increased body size than does the smaller sex. A consequence of this is that we might expect the larger sex to show a greater response to environmental factors that enhance body size. This heightened sensitivity is referred to as the condition dependence hypothesis. Condition dependence is a form of phenotypic plasticity whereby the expression of a trait changes with the phenotypic condition of the individual [11]. The condition dependence hypothesis was originally developed to explain the evolution of sexually-selected traits that signal mate quality in a reliable, condition-dependent manner [11–15]. However, the condition dependent hypothesis can be applied more generally to any trait that is differentially selected in males and females, regardless of the mode of selection [6]. If individuals in good condition enjoy larger marginal benefits from increasing trait size in one sex relative to the other, this sex should show greater investment in the trait, be it overall body size or the size of an individual organ [16,17]. In as much as condition reflects environmental variation, particularly access to nutrition during growth, the condition dependent hypothesis predicts a positive correlation between SSD and size plasticity such that the larger sex will exhibit greater size plasticity than the smaller sex [18]. From a quantitative genetic perspective, this sex-specific plasticity (SSP) can be considered a *GxE* interaction, where *G* is sex and *E* is environmental variation.

Several studies provide evidence for a correlation between sexual dimorphism and sex-specific plasticity, either by comparing multiple dimorphic and non-dimorphic traits within species, or by comparing the same trait across multiple species. For example, the strength of condition dependence increases with the degree of sexual dimorphism among different traits in *Prochyliza xanthostoma* and *Telostylinus angusticollis* flies [12,19]: traits that are more dimorphic show greater differential plasticity between sexes. The same pattern is observed for sexually-dimorphic traits among species [20]. For the specific example of SSD, where the trait is body size as a whole, comparative studies among species also support the condition dependent hypothesis. In arthropods, where SSD is female-biased, there is a general trend for female body size to be more plastic[18,21]. Among birds and mammals, where SSD is typically male-biased, the opposite is true [22,23]. Further, among the few arthropod taxa with male-biased SSD, males body size also appears to be more plastic than female body size [6].

What is unclear from these studies, however, is the origin of these correlations. One possibility is that SSP evolves as a consequence of SSD, with selection targeting different developmental-genetic mechanisms for each. Indeed, early theoretical models of the condition dependent hypothesis argue that sexually dimorphic traits are more likely to evolve if the loci that regulate expression of the traits are distinct from but linked to loci that cause the traits to be condition-dependent [11].

Alternatively, selection for SSD may target the same developmental-genetic mechanisms that increase size under better conditions, that is, selection may focus on the mechanisms that regulate size plasticity. From this perspective, SSD evolves as a consequence of sex-specific responses to changes in condition. Distinguishing between these hypotheses requires insight into whether SSD and SSP covary among individual genotypes within a population, which is not provided by existing data.

Here we investigate whether SSD and SSP are genetically correlated among individuals within a population by manipulating nutrition and measuring body size and body-size plasticity in 196 isogenic *D. melanogaster* lineages. We found considerable genetic variation for SSD among genotypes, and determined that variation in SSD and SSP are genetically correlated. These data support the hypothesis that same loci regulate both SSD and SSP and suggest that the observed SSD is a consequence of selection for SSP.

## Results

### Sexual Size Dimorphism shows genetic variation in Drosophila melanogaster

We found significant genetic variation in SSD among the 186 DGRP lineages that we measured (**Fig. 1**), using pupal area of fed flies as a measure of overall body size [24]. Specifically, a linear mixed effect model that included random variation in SSD among lineages was a significantly better fit to the data than a model that did not (**Table 1**). Further, SSD is female biased (Best Linear Unbiased Predictions: male body size = 14.47, female body size = 14.56).

**Table 1:** Effect of including random variation in sex-by-lineage interaction on the fit of the relationship between body size and sex.

| Model<sup>a</sup> | DF<sup>b</sup> | AIC<sup>c</sup> | BIC<sup>c</sup> | Likelihood Ratio<sup>c</sup> | P<sup>d</sup> |
| --- | --- | --- | --- | --- | --- |
| Model 1: $P = S + B + S \cdot L$ | 7 | -11764 | -11715 | 40.2 | <0.001 |
| Model 2: $P = S + B + L$ | 5 | -11728 | -11693 | | |

<sup>a</sup> P is the log transformed pupal case area, S is the sex, B is block (random effect), L is the lineage (random effect). The models differ by having random sex-by-lineage effects (model 1) versus random lineage effects (model 2).
<sup>b</sup> Estimated degrees of freedom for each model.
<sup>c</sup> AIC, BIC, and likelihood ratio calculated using ML fit.
<sup>d</sup> P-value calculated by parametric bootstrapping using ML fit.

**Figure 1:**
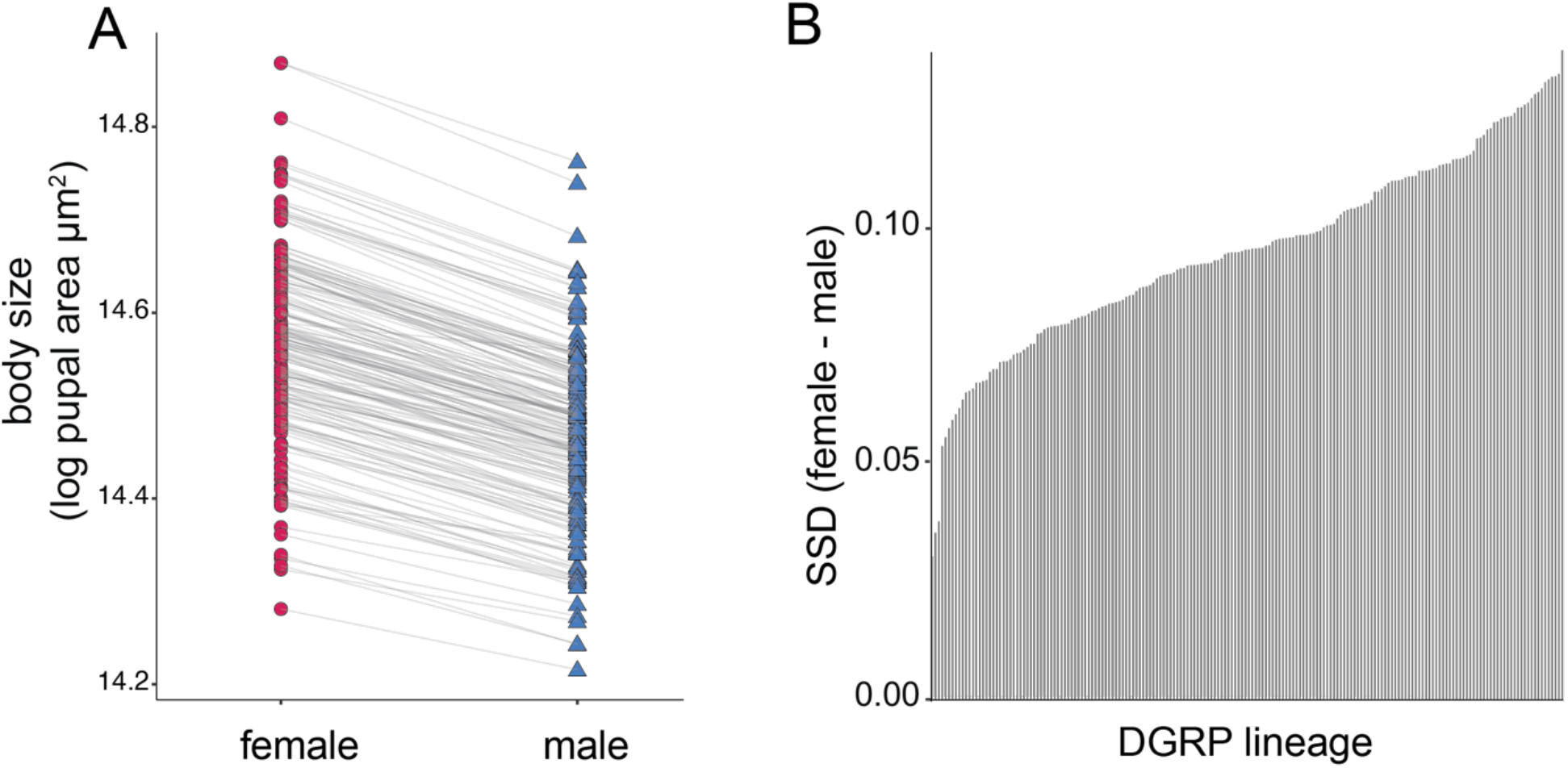
Variation in Sexual Size Dimorphism (SSD) among isogenic lineages. (A) Female and male body size for each of the 186 lineages. Estimates of body size are Best Linear Unbiased Predictions (BLUPs) from a mixed model linear regression (Table 1: Model 1). Sexes for each lineage are connected by a line. (B) Ranked plot of SSD for these lineages (*log body size*_*female*_ *– log body size*_*male*_).

### Variation in SSD reflects variation in female body size more than male body size

To determine the degree to which SSD results from variation in male versus female size, we fit linear relationships between SSD (*y*) and either male or female body size (*x*) (**Fig. 2A**). If variation in SSD is driven by variation in female size more than variation in male size, the slope of this relationship should be steeper for females than for males, and the *R*^*2*^ should be higher. We found that the slope of the relationship between body size and SSD was significantly steeper for females than males (t-test on the slope, *P* < 2e-16), and *R*^*2*^ was also significantly higher for female size than male size against SSD (R^2^(female)= 0.29, 95% CI: 0.18 - 0.39; R^2^(male)= 0.14, 95% CI: 0.05 - 0.23), **Fig. 2A**). These data indicate that variation in SSD among lineages was due primarily to variation in female size.

**Figure 2:**
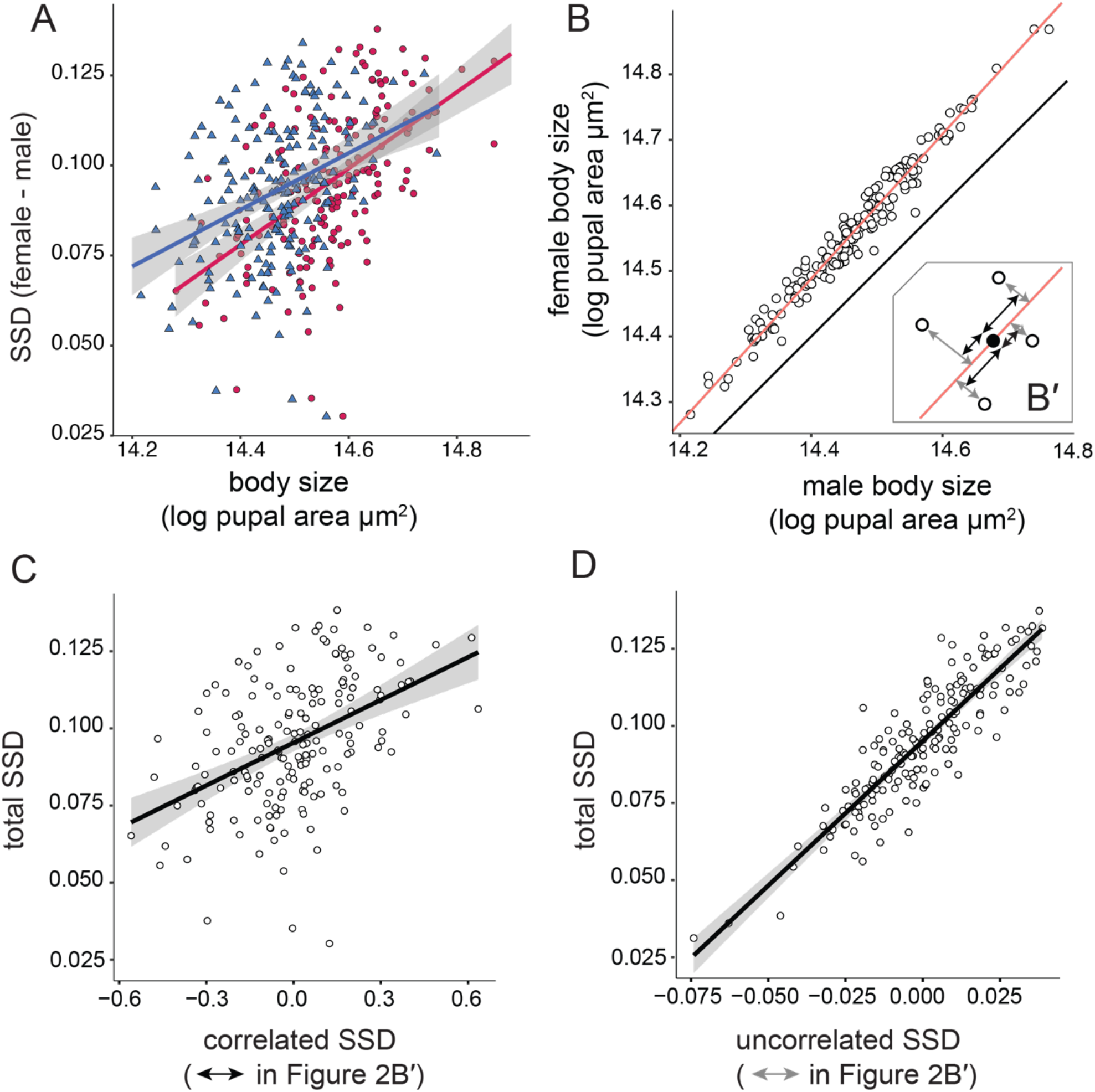
Patterns of SSD variation among lineages. (A) OLS linear regression between female (red circle) and male (blue triangle) size and SSD. Variation in SSD is due more to variation in female (*R*^*2*^ = 0.29, 95% CI: 0.18 - 0.40) than male (*R*^*2*^ = 0.29, 95% CI: 0.18 - 0.40) size. (B) Major axis regression of female on male size (red line, slope = 1.1). SSD increases as lineage pupal size increases, although more so in females than in males. Black line is *x=y*. (B ’: Inset) Variation in SSD can be explained by variation that is correlated with sex-averaged lineage size (correlated SSD, black arrows) and variation that is uncorrelated with sex-averaged lineage size (uncorrelated SSD, gray arrows). Solid point represents the bivariate mean female and male body size among all lineages. (C & D) OLS linear regression of total SSD against correlated SSD (C) and uncorrelated SSD (D). Correlated SSD accounts for 22% variation in variation in SSD among lineages (*R*^2^ = 0.22) whereas uncorrelated SSD accounts for 78% variation in variation in SSD among lineages (*R*^2^ = 0.78). Gray shading is 95% confidence interval of the fit.

### SSD increases with sex-averaged body size among lineages

Male and female size are tightly correlated among lineages (Pearson’s correlation R = 0.98, p<2.2e-16, **Fig. 2B**). If variation in SSD due is more to variation in female size than male size, it follows that regressing female size (y-axis) against male size (x-axis) should yield a slope significantly greater than 1. Major Axis (MA) regression between female and male body size supported this prediction (slope= 1.10, 95% CI = 1.07-1.13; **Fig. 2B**). The observation that females lie above the line *y=x* (**Fig. 2B**) indicates that females are larger than males among the lineages. The observation that the slope of this line is > 1 indicates that SSD increases with sex-averaged body size among lineages (OLS, *P=* 8.062e-09, **Fig. S1**) due to a disproportionate increase in female size. (Here, sex-averaged body size refers to body size independent of sex, that is 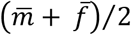, where 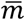 and 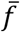 are predicted male and female body size respectively). Importantly, this suggests that there are two factors that generate variation in SSD among lineages: Variation in SSD that is correlated with variation in sex-averaged body size among lineages, and variation in SSD that is uncorrelated with variation in sex-averaged body size. We refer to these effects as *correlated-SSD* and *uncorrelated-SSD*, respectively. In **Figure 2B’**, correlated-SSD is the variation in SSD generated as lineages move up or down the red line (black arrows), while uncorrelated-SSD is the variation in SSD generated by lineage lying above or below the red line (grey arrows).

To calculate the extent to which variation in SSD is due to effects that are correlated and uncorrelated with sex-averaged body size, we first extracted the fitted values and residuals for each lineage from the MA regression of female on male body size. The *R*^*2*^ of the linear relationship between the fitted values and SSD captures the proportion of SSD variation that is due to variation in sex-averaged body size (correlated-SSD), while the *R*^*2*^ of the linear relationship between the residuals and SSD captures the proportion of SSD variation that is independent of variation in sex-averaged body size (uncorrelated-SSD). Because total variation in SSD is a combination of both variation that is correlated and uncorrelated with variation in sex-averaged body size among lineages, the adjusted *R*^2^ of these two relationships sum to one. We found that the *R*^*2*^ for the sex-averaged dependent effect was 0.1832 (95% CI: 0.12–0.32) (**Fig. 2C**), while the *R*^*2*^ for the sex-averaged independent effect was 0.8107 (95% CI: 0.73–0.84) (**Fig. 2D**). Consequently, while variation in sex-averaged body size among lineages partially explains variation in SSD, variation due to difference in male or female size independent of sex-averaged body size explains variation in SSD to a greater extent.

### Females have greater nutritionally-induced size plasticity than males

We first explored the factors that generate variation in SSD that are correlated with variation in sex-averaged size among lineages (correlated-SSD). Because female size is more variable among lineages than is male size, it follows that females are more sensitive to factors that influence size than are males. A key environmental determinant of body size in all animals is nutrition during growth and development. We therefore tested whether females are more sensitive than males to variation in developmental nutrition, using the difference in average male or female size between fed and starved treatments as a measure of the male or female plasticity for each lineage. In nearly all of the DGRP lineages, females had higher nutritionally-induced size plasticity than males (**Fig. 3A & B**), and there was significant interaction between the effects of sex and starvation treatment on body size across lineages (Table 2, Model 3: *P*_sex*starvation treatment_ < 0.0001), indicating sex-specific plasticity. If females are generally more sensitive to size-regulatory factors than males, whether those factors are environmental or genetic, then this increased sensitivity may account for variation in SSD that is correlated with the sex-averaged body size of a lineage.

**Table 2:** Effect of including random variation in sex-by-starvation-by-lineage interaction on the fit of the relationship between body size, sex, and starvation treatment.

| Model<sup>a</sup> | DF<sup>b</sup> | AIC<sup>c</sup> | BIC<sup>c</sup> | Likelihood Ratio<sup>c</sup> | P<sup>d</sup> |
| --- | --- | --- | --- | --- | --- |
| Model 3: $P = S \cdot D + B + S \cdot D \cdot L$ | 16 | -19252 | -19137 | 15.11 | 0.004 |
| Model 4: $P = S \cdot D + B + (S+D) \cdot L$ | 12 | -19259 | -19160 | | |

<sup>a</sup> P is the log transformed pupal case area, S is the sex, D is the number of days of starvation, B is the collecting block (random effect), L is the lineage (random effect). The models differ by having random variation in SSP among lineages (sex-by-starvation-by-lineage interaction, model 3) versus random additive effect of sex and starvation among lineages.
<sup>b</sup> Estimated degrees of freedom for each model.
<sup>c</sup> AIC, BIC, and likelihood ratio calculated using ML fit.
<sup>d</sup> P-value calculated by parametric bootstrapping using ML fit.

**Figure 3.**
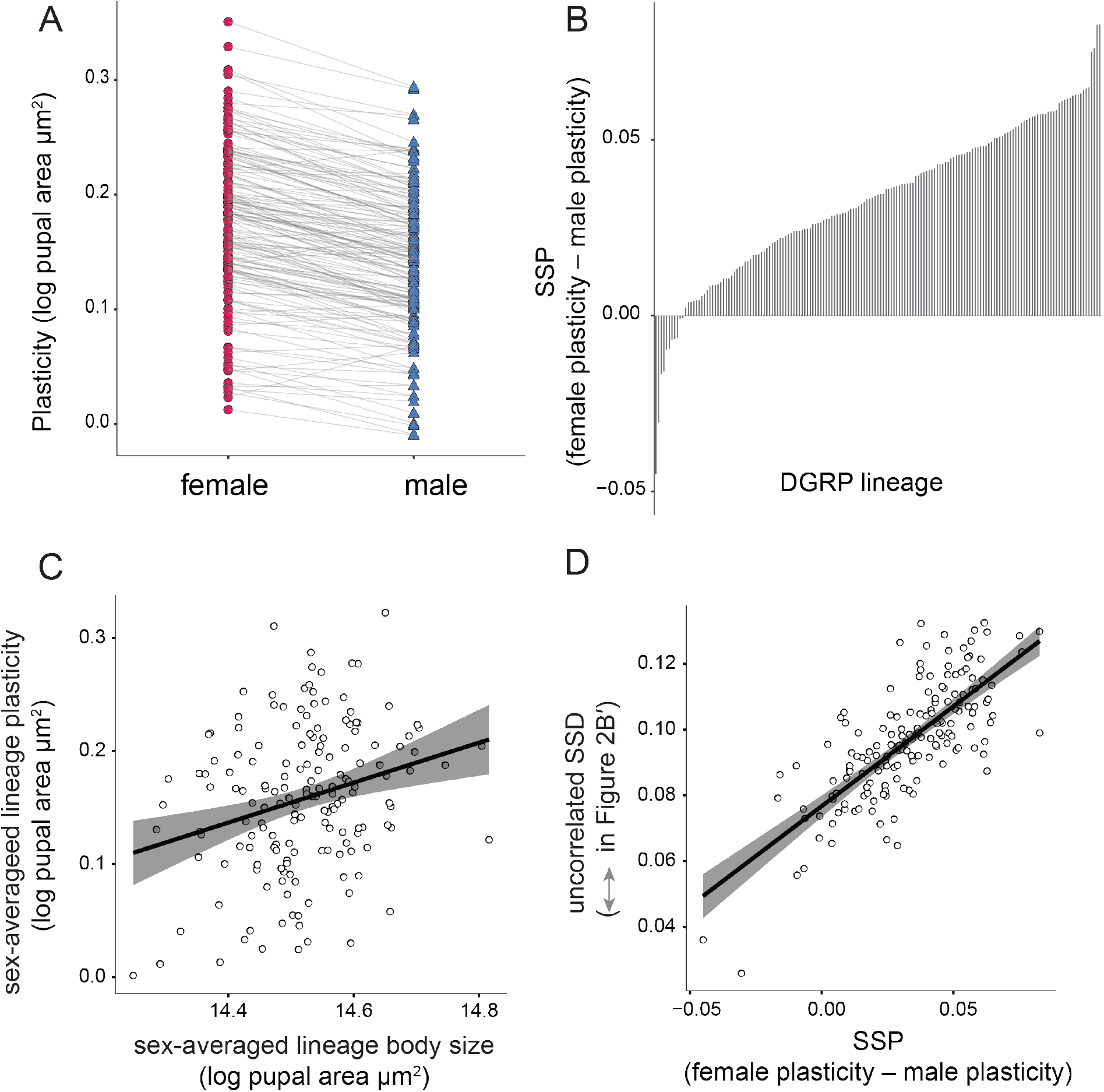
Patterns of SSP variation among lineages. (A) Female and male size plasticity for each of the 186 lineages. Estimates of plasticity use Best Linear Unbiased Predictions (BLUPs) of body size from a mixed model linear regression on fed and starved flies (Table 1: Model 1).. (B) Ranking plot of SSP for these lineages, where SSP is female plasticity – male plasticity (C) OLS linear regression of sex-averaged lineage body size against sex-averaged lineage plasticity. The relationship is significant but lose (*R*^2^ = 0.07, *P* = 0.0005). (D) OLS linear regression between uncorrelated SSD and SPP among lineages. The relationship is strong and highly significant. (*R*^2^=0.56, *P* <2.2e-16). Gray shading is 95% confidence interval of the fit.

### Sex Specific Plasticity covaries with Sexual Size Dimorphism independent of lineage body size

One simple hypothesis as to why females are more plastic than males is that females are larger than males and larger individuals are more plastic than smaller individuals. Indeed, there is a significant relationship between sex-averaged size and sex-averaged size plasticity among lineages (**Fig 3C**). However, while significant, this relationship is weak (*R*^2^ = 0.07, *P* = 0.0005), suggesting that SSP is not simply a reflection of the fact that females are bigger than males. To address this further, we explored how variation in SSP covaries with SSD independently of sex-averaged size, i.e. uncorrelated SSD (**Fig. 2D**). We first tested whether SSP varied significantly among lineages, and found that it did: A linear mixed effect model that included random variation in SSP (sex and starvation interaction) among lineages was a significantly better fit to the data than one that did not (Table 2). (Best Linear Unbiased Predictions: fed male body size = 14.47, fed female body size = 14.56, starved male body size = 14.42, starved female body size = 14.39).

We then whether variation in SSP among lineages correlates with the (1) overall variation in SSD among lineages, and (2) variation in SSD independent of sex-averaged size (uncorrelated-SSD). In as much as variation in SSD among isogenic lineages is due to genotype, there was a positive genetic correlation between SSD and SSP, consistent with the condition dependent hypothesis of SSD (**Fig .3D**; Pearson’s *R* = 0.75). Further, an OLS linear regression of SSP against uncorrelated-SSD was significant with an adjusted *R*^2^ of 0.56. Thus variation in SSP among lineages explains the majority of variation in SSD that is independent of sex-averaged lineage pupal size.

## Discussion

Sexual size dimorphism is widespread in animals. Among the possible evolutionary mechanisms that generate sex-specific size differences, the condition dependence hypothesis proposes that when the larger sex experiences stronger directional selection on body size, that sex will have greater fitness benefits if it can exploit environmental conditions favourable to growth [14,16,17], resulting in sex-specific size plasticity [12,19,20,25]. Numerous comparative studies demonstrate a correlation between SSD and sex-specific size plasticity among traits within species [12], among closely related species [18,20,26], and among populations within species [8,27]. Perhaps because of the correlative nature of these data, the condition dependence hypothesis is often discussed as a process that shapes SSD rather than one that generates it in the first place. Sex specific plasticity, however, *de facto* generates SSD, and so the condition dependent hypothesis can explain the evolution of SSD if selection for SSD targets the same developmental-genetic mechanisms that generate SSP. This predicts a positive genetic correlation between variation for SSD and SSP among individuals within a population. This is problematic to test, however, because SSD is a population-level, not an individual-level, measure. To circumvent this complication, we estimated SSD, SSP, and their covariation from populations of isogenic *Drosophila* lineages, where body size variation within lineages was induced by manipulating access to food. We found significant variation in female-biased SSD among the lineages, and a tight positive correlation between SSD and female-biased SSP. That is, in genotypes where females are larger relative to males and SSD is more pronounced, independent of the sex-averaged body size of the lineage, females are correspondingly more plastic. The genetic covariation between SSD and SSP is consistent with the hypothesis that both are regulated by the same developmental mechanisms, and this pleiotropy has important implications for origin and evolution of SSD, which we explore below.

### Proximate mechanisms for Sex Specific Plasticity and Sexual Size Dimorphism

Previous studies have used a variety of data to test the condition dependent hypothesis, correlating SSD with SSP among multiple traits within species [12,28], or correlating SSD with SSP for a single trait among populations [8,27] or species [20,21]. However, while these correlational studies provide general support for the condition dependence hypothesis, they cannot determine if condition dependence of sexually-dimorphic traits arise as a consequence of the dimorphism, or if condition dependence generates sexual dimorphism in plastic traits [16]. In order to distinguish between these two hypotheses it is necessary to have some understanding of the developmental genetic mechanisms that regulate both SSD and SSP. Our study goes some way to addressing this, by demonstrating a tight genetic correlation between SSD and SSP. There are two major causes of genetic correlation: linkage disequilibrium (LD) or pleiotropy. The DGRP population of *D. melanogaster* in our study has a rapid decay of LD, with an average squared correlation below 0.2 at just 10 base pairs on the autosomes [29]. Consequently, it seems unlikely that the genetic correlation we observe between SSD and SSP among lineages is the result of linkage.

A number of recent studies support the hypothesis that SSD in *Drosophila* is generated by the same developmental-genetic mechanisms that regulate nutritional plasticity of body size; specifically the insulin/IGF-signalling (IIS) and TOR-signalling pathways (see [30–32]. The IIS responds to circulating insulin-like peptides (dILPS in *Drosophila*), which are released in a nutrient-dependent manner and bind to the insulin receptors (InR) of dividing cells [33]. Binding activates a signal-transduction pathway that ultimately controls the expression of genes involved in regulating cell survival, growth, and proliferation [34]. The TOR-signalling pathway has a similar effect, but is regulated more directly by circulating amino acids [32,35]. There is considerable cross-talk between the two pathways so that they work together to tune organismal growth and development in response to the nutritional environment [36].

In *Drosophila*, SSD is eliminated in flies carrying a hypomorphic mutation of *InR* [37], which indicates that a functional IIS-pathway is necessary to generate the difference in body size between males and females. Further, fat body-specific knock-down of the sex-determining gene *Transformer* – which produces protein in females but not in males – results in the retention of dILP2 in the insulin-producing cells in the brains of females but not males [38]. Retention of dILP2 correlates with lower circulating levels of dILP2, and these females have a correspondingly reduced body size. These data suggest that females are larger than males because they have higher levels of circulating dILPs when well fed. Subsequent work has demonstrated that females, but not males, increase body size when reared on a very-high protein diet, and that this elevated sex-specific plasticity is also dependent on dILP2 and Transformer [39]. Collectively, these data support the hypothesis that both SSD and SSP are regulated by the IIS-pathway in *Drosophila*. An added nuance, however, is that the effect of nutrition on body size depends not just on food quantity but also food quality. Specifically, females, but not males, reduce body size when reared on high sugar diets ([40]although see [39]), and this appears to reflect sex-specific differences in the activity of the TOR-signalling pathway [41,42].

Sexual dimorphism in the size of sexually-dimorphic traits in other systems also appears to be regulated by the IIS pathway. The horns of male rhinoceros beetle (*Trypoxylus dichotomus*) show heightened insulin-sensitivity and corresponding nutritional sensitivity relative to other traits in the body, with knock-down of *InR* having a greater effect on male horn size - reducing the sexual dimorphism [43]. Similarly, the size of the exaggerated mandibles of male broad-horned flour beetle (*Gnatocerus cornutus*) is regulated by a specific ILP (GcorILP2), which is expressed in a condition-dependent manner [44]. Thus, increased activity of the insulin-signaling pathway, either systemically or in specific tissues, appears to account for sex-specific condition-dependent increases in trait size among a diversity of insects.

### Evolutionary implications of pleiotropy for SSP and SSD

The observation that sexual size dimorphism in *Drosophila* is generated by sex-specific plasticity has important implications for our understanding of the evolution of sexual dimorphism in general and SSD specifically. In particular, it suggests that selection targets mechanisms that conditionally increase trait size in one sex over the other. Why the conditional increase in trait size is favored only in one sex depends on the mode of selection. For sexually-dimorphic traits that are used by males to attract or compete for females and are subject to sexual selection, heightened conditionality of trait expression ensures that the trait is a reliable and sensitive indicator of male quality[43]. In flies and other insects, however, SSD appears to a consequence of fecundity selection. Female reproductive success is limited by the number of eggs she lays, and fecundity increases with body size [39,45–47].

Further, artificial selection on fecundity in female *Drosophila* leads to an increase in SSD [48]. The effect of male body size on fitness in *Drosophila* is much weaker or undetectable [39,45], although this may be because male mating success is limited by access to females, and studies exploring the effect of male body size on fecundity used single pair matings [39,45]. While larger males may have slight reproductive advantage under scramble competition for matings [49], it is possible that males on high quality diets utilize mechanisms other than body size to enhance fitness, for example by reducing developmental time or increasing the number of matings. Regardless of the form or target of selection, however, selection favouring SSP in size will necessarily generate SSD, and our data suggest that selection targets the same mechanisms to result in both..

Our data showing a genetic correlation between SSD and SSP also has interesting implications for Rensch’s rule, which states that male-biased SSD increases with overall body size, while female-biased SSD decreases with overall body size, among species or populations [5]. The general explanation for this phenomenon is that male body size varies more than female body size among populations and species, a pattern that has been hypothesized to result from greater plasticity in male size [50]. Interspecific variation in SSD consistent with Rensch’s rule is common in vertebrates [51,52], but is also seen in several invertebrate groups, including the Drosophilidae [53]. Among *Drosophila* genotypes, however, we found female size to be more variable than male size, and so female-biased SSD increases with overall body size among lineages - the reverse of Rensch’s rule (albeit applied to genotypes rather than populations or species). The observation that the generally female-biased interspecific variation in SSD among Drosophilidae follows Rensch’s rule, but that intraspecific variation in SSD exhibits a reverse Rensch’s rule, suggests that Rensch’s rule cannot be explained by greater plasticity in males versus females. Indeed, there is a general trend in both vertebrates and invertebrates that the larger sex is more plastic than the smaller sex [18,22]. If Rench’s rule were due to sex-specific plasticity, then we should observe Rensch’s rule only when males are larger than females, but not when females are larger than males.

## Conclusions

We tested whether the condition dependence hypothesis explains sexual size dimorphism using individual variation in a population of *Drosophila melanogaster*. We found that SSD and SSP shows a strong positive correlation, suggesting shared developmental-genetic mechanisms. We further suggest that the selective pressures that generate the SSD act on the same developmental mechanisms that regulate nutritionally-induced body size plasticity. In other words, the observed sexual-size dimorphism may be a consequence of sex-specific selection favouring increased body size as condition improves in females but less so in males. Full testing of this hypothesis requires identification of the loci that underlie variation in both SSD and SSP among these *Drosophila* lineages. More generally, however, our study makes it clear that to fully understand that selective pressures that generate SSD, it is necessary to also understand that developmental-genetic mechanisms those selective pressures target.

## Methods

### Fly stocks

Flies come from the *Drosophila* Genetic Reference Panel (DGRP), consisting of more than 200 inbred *Drosophila melanogaster* lineages generated from a natural population in Raleigh, NC, USA, after 20 generation of full-sib mating [29]. Each lineage discussed in this paper refers to a specific isogenic line in the DGRP with a unique genotype.

### Starvation treatment

For each DGRP lineage, flies were reared following previously publish protocols [24,54,55]. Eggs were collected over 12-20 h and transferred in lots of 50 into 7ml fly food, and the larvae were reared at 22ºC, in on standard cornmeal-molasses medium until the starvation treatment was applied. Larvae were removed from food at precisely timed developmental stages and starved for either 0–24 h or 24-48h before pupation to generate variation in body size. Because larvae stop feeding approximately 24h before pupation [56], larvae removed from the food 0–24 h before pupation were essentially allowed to feed *ad libitum* and are referred to as ‘fed’ flies. In contrast, larvae removed from the food 24-48h before pupation were starved during the period when adult body size is affected by nutrition [57], and are referred to as ‘starved’ flies. After being removed from the food, larvae were transferred to an empty vial containing a wet cotton plug and left until pupariation. Pupae were then transferred to individual 2.5 mL Eppendorf tubes until eclosion to allow us to associate each adult fly with its pupal case. After the flies emerged as adults from their pupal cases, they were sexed and the pupal case was imaged and size quantified using semi-automated software developed in the Shingleton lab [58]. We use area of the pupal silhouette (dorsal view) as a proxy for adult body size [24,54,55]. Flies were collected in nine temporal blocks, with one lineage repeated across all blocks to serve as a control.

### Statistical analyses

All analyses were conducted in R (v. 4.0.3), and the script and data used in the analyses provided on Dryad. Prior to analysis, data were filtered such that each sex-treatment-lineage combination had at least 10 pupal area measurements. After filtering, the remaining DGRP lineages were analyzed. All measurements were log transformed prior to analysis.

We used a Mixed Linear Model (R package: *lme4*) [59] to test for genetic variation in Sexual Size Dimorphism (SSD) and Sex Specific Plasticity (SSP).

For SSD we used only the data from the fed flies and fit the models:

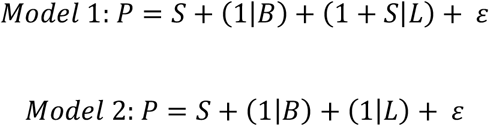

where *P* is body size, *S* is sex, *B* is block (random factor), *L* is lineage (random factor), and ε is error. The two models differ only in whether there is random variation in the effect of sex on body size among lineages (Model 1), or not (Model 2); that is whether SSD varies among lineages. We compared the models using a log likelihood ratio test and parametric bootstrapping.

For SSP, we used the data from the fed and starved flies and fit the models:

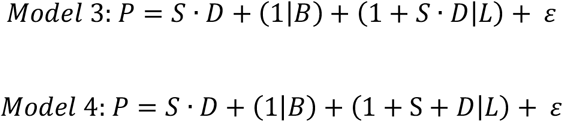

where *D* is diet (fed or starved). The two models differ in whether there is random variation in the interaction between the effects of sex and diet on pupal size among lineages (Model 1), or not (Model 2); that is whether SSP varies among lineages. Again, we compared the models using a log likelihood ratio test and parametric bootstrapping.

For the analysis of genetic variation of SSD, we used the Best Linear Unbiased Prediction (BLUP) values for each lineage and sex from Table 1: Model 1 to estimate predicted male and female size (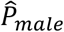 and 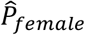 respectively) for each lineage for both fed and starved flies separately.

Predicted sex-adjusted body size for fed flies in each lineage was calculated as:

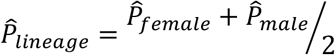

Sexual size dimorphism for a lineage was calculated as:

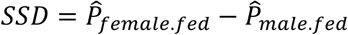

Nutritionally-induced size plasticity for each sex within a lineage was calculated as:

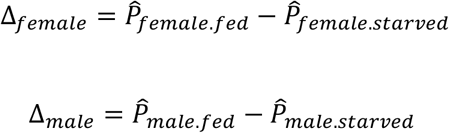

Mean sex-adjusted size plasticity for each lineage was calculated as:

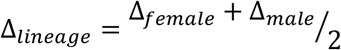

Sex-specific size plasticity for a lineage was calculated as:

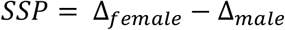

These values were then used to examine the relationship between female and male size using Major Axis (MA) regression (R package: smartr) [60], and the fitted and residual values for this relationship were used to analyze sex-averaged size dependent and sex-averaged size independent measures of SSD (referred to as correlated and uncorrelated SSD, respectively). We then used ordinary least squares (OLS) regression to explore the relationship between total SSD and uncorrelated and correlated SSD, lineage mean size and lineage mean plasticity, and between SSP and correlated and uncorrelated SSD.

## Acknowledgements

We would like to thank all the members of the Shingleton and Frankino labs for their assistance and support during the collection of these data. Particular thanks to Omid Ziabari, Sonya Messer and Chris Edomwande for organizing the collection of the pupal measurements.

---

**Supplementary Figure 1:**
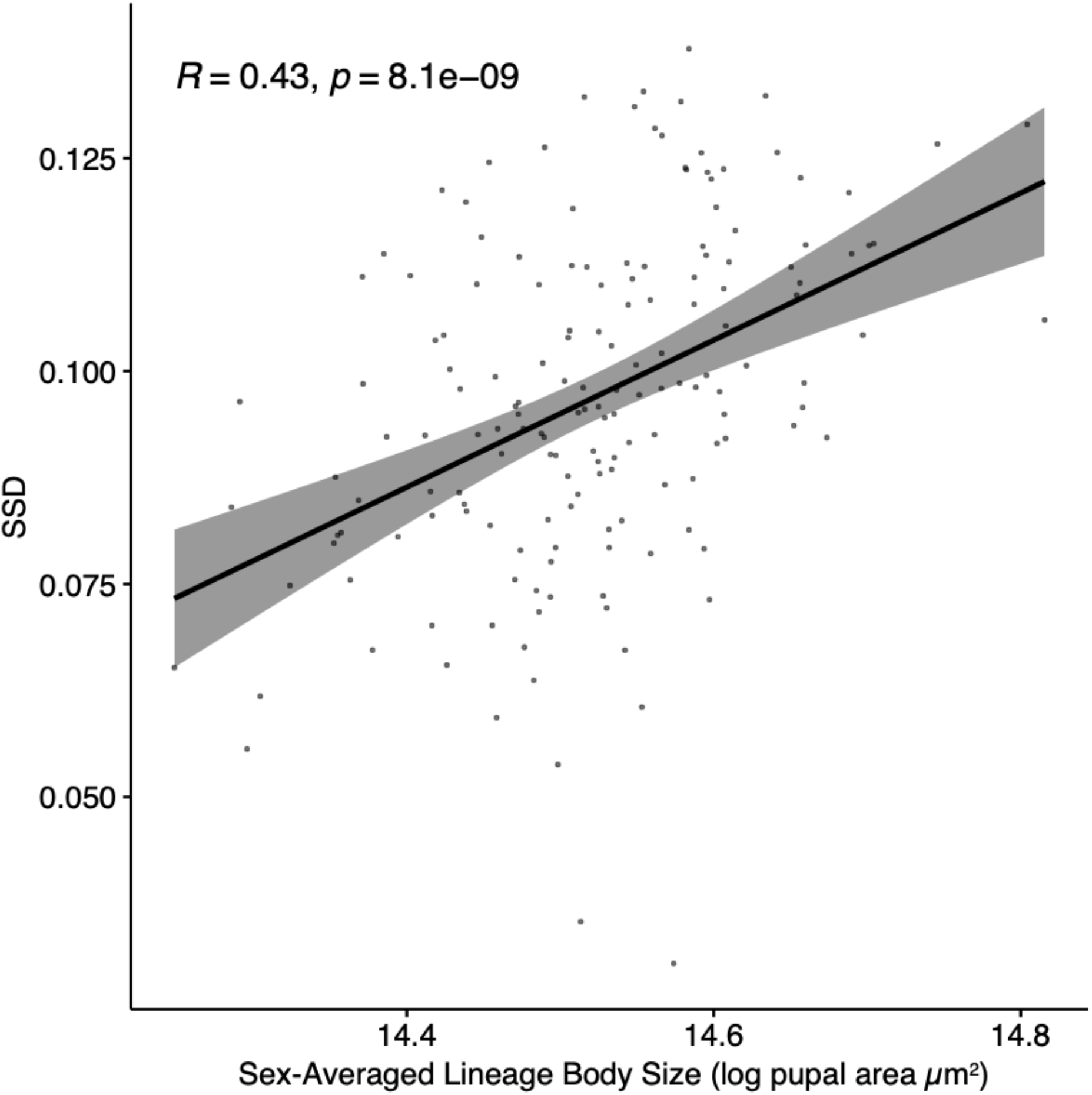
The relationship between sex-averaged lineage body size and SSD. Line is an OLS linear regression. Shading is 95% confidence interval of the slope.

